# The Effects of Preeclamptic Milieu on Cord Blood Derived Endothelial Colony-Forming Cells

**DOI:** 10.1101/2023.12.03.569585

**Authors:** Eva Hall, Laura Alderfer, Erin Neu, Sanjoy Saha, Ellie Johandes, David M. Haas, Laura S. Haneline, Donny Hanjaya-Putra

## Abstract

Preeclampsia is one of the leading causes of infant and maternal mortality worldwide. Many infants born from preeclamptic pregnancies are born prematurely with higher risk of developing cardiovascular later in their life. A key mechanism by which these complications occur is through stress-induced dysfunction of endothelial progenitor cells (EPCs), including endothelial colony-forming cells (ECFCs). To gain insight into this, cord blood derived ECFCs isolated from preeclamptic pregnancies (PRECs) were analyzed and compared to their healthy counterparts. While PRECs preserve key endothelial markers, they upregulate several markers associated with oxidative stress and inflammatory response. Compared to ECFCs, PRECs also exhibit lower migratory behaviors and impaired angiogenic potential. Interestingly, treatment of neuropilin-1 can improve tube formation *in vitro*. Collectively, this study reports that preeclamptic milieu influence phenotypes and functionality of PRECs, which can be rejuvenated using exogenous molecules. Promising results from this study warrant future investigations on the prospect of the rejuvenated PRECs to improve lung function of infants born from preeclamptic pregnancies.

## Introduction

Every year, preeclampsia, a hypertensive disorder of pregnancy seen after 20 weeks of gestation, affects up to 8% of pregnancies and causes approximately 500,000 fetal and 70,000 maternal deaths making it one of the leading causes of maternal and infant death worldwide^1–3^. Despite it being a dangerous disease, there are very few treatments. Currently, many methods focus on treating maternal symptoms, including increased blood pressure and seizures, and monitoring fetal development^4,5^. However, the only definitive treatment is delivery, but even this is not a perfect solution^1,6^. There are outstanding, long-term risks for both the mother and the child including increased risk of cardiovascular disease, stroke, and hypertension even post-delivery^2^. Additionally, preterm delivery can cause a higher incidence of systemic complications in the infant such as diabetes, neurodevelopmental impairment, and bronchopulmonary dysplasia (BPD)^3,7,8^.

One of the challenges with treating preeclampsia is the root cause is not fully understood as it a complex, multisystem pathology. Due to this, it is often referred to as a “disease of theories”^9,10^. There are some mechanisms which now are known to play a role in the pathology including perturbations in the renin-angiotensin system, excessive immune responses, oxidative stress, and other various genetic factors, but no overarching hypothesis has been developed^11–19^. Endothelial dysfunction, especially during spiral remodeling, and imbalanced angiogenic factors are also shown to play a major role. During a normal pregnancy, the spiral arteries in the placenta are remodeled by trophoblasts, which converts them to an invasive phenotype and allows them to enter the smooth muscles of the uterus known as the myometrium. These vessels remodel into high capacitance, high flow vessels due to the loss of endothelium and muscle. However, in preeclampsia the arteries do not become invasive or remodel, which can lead to ischemia and hypoxia^2,3^.

Some angiogenic factors including vascular endothelial growth factor (VEGF), placental growth factor (PlGF), and transforming growth factor (TGF-β1) are also dysregulated in preeclampsia and are part of the reason for the spiral artery invasion failure. These growth factors recruit circulating endothelial progenitor cells (EPCs) and stimulate vascularization of the placenta in a healthy pregnancy^20,21^. However, in preeclampsia an increase in soluble fms-like tyrosine kinase-1 (sFlt-1), also known as soluble VEGFR-1, binds to VEGF and PlGF. This prevents these growth factors from providing their normal angiogenic affects and can prevent spiral artery remodeling^2,22–24^. While soluble endoglin (sEng), disrupts TGF-β1 signaling, which in turn prevents spiral artery invasion^25^.

EPCs, especially endothelial colony forming cells (ECFCs), play a major role in maintaining vascular homeostasis and can help form *de novo* vasculature^26^. ECFCs are highly proliferative and can form vessels *in vivo*^27,28^. In addition, they have higher angiogenic potential than some mature endothelial cells such as human umbilical vein endothelial cells (HUVECs)^29^. *In utero* diseases have been seen to influence ECFCs, which in turn affects proliferation, migration, and tube formation^30–32^. However, healthy ECFCs have also seen to have therapeutic potential to mitigate some of the effects of these diseases, including assisting with chronic lung disease and neurovascular repair in growth restricted limbs^33–35^.

In this study, we report that cord blood derived ECFCs isolated from preeclamptic pregnancies (PRECs) have different functional and genetic characteristics compared to their healthy counterparts. Proliferation and senescence are increased, while migration and tube forming ability are decreased. The cytokines and angiogenesis associated genes from PRECs are dysregulated compared to healthy ECFCs. Interestingly, there is also an increase in lymphatic markers in the PRECs. Collectively, we demonstrate PRECs exhibit impaired functional and genetic characteristics, which can be rescued using various soluble factors.

## Methods

### Human ECFCs

The human ECFCs were collected from cord blood from pregnancies with a variety of gestational ages **(Table 1)** and provided by members of the Indiana University School of Medicine, Dr. Laura Haneline and Dr. David Haas. They were cultured in Endothelial Cell Growth Medium 2 (EGM2; PromoCell) and were used between passage number 4 and 8, except for when testing for senescence when passage number 12 was used. The ECFCs were incubated at 37°C with 5% CO2 and were passaged at 70%-90% confluency. All cells were seeded onto 50µg/mL Rat Tail Collagen 1 (RTC1; Corning). All cell lines were routinely tested for mycoplasma contamination and were negative throughout this study.

**Table 1.** ECFC sample characteristics and their associated clinical data.

| Group | Sample ID | Gestational age at birth (weeks) | Baby's Gender | 50g 1 hour GTT |
| --- | --- | --- | --- | --- |
| Control | E1-CB-111 | 38 | Male | 102 |
| Control | E1-CB-130 | 41 | Female | 116 |
| Control | E1-CB-150 | 39 | Male | 115 |
| Control | E1-CB-153 | 39 | Male | 115 |
| Control | E1-CB-157 | 39 | Female | 139 |
| Preeclampsia | PREC 348 | 36 + 1 | Male | 114 |
| Preeclampsia | PREC 370 | 37 + 2 | Male | 109 |
| Preeclampsia | PREC 304 | 37 + 0 | Female | 118 |
| Preeclampsia | PREC 349 | 38 + 6 | Male | 108 |
| Preeclampsia | PREC 367 | 31 + 3 | Male | 112 |
| Preeclampsia | PREC 340 | 39 + 0 | Female | 120 |
| GDM | E1-CB-74 | 39 | Male | 239 |

### Flow Cytometry

Cells were grown until they reached approximately 80%-90% confluency in a T75 flask (VWR). They were harvested and diluted to 1x10^6^ cell/mL in FACS Buffer (Thermo Fisher). 100 µL of cell suspension were added to wells of a 96-well round bottomed plate (Thermo Fisher). APC Conjugated CD31 (R&D Systems), PE Conjugated CD34 (Abcam), PE Conjugated CD144 (R&D Systems), and APC Conjugated NRP-1 (R&D Systems) antibodies were added at the concentration or dilution recommended by their manufacturers and allowed to incubate for at least 30 minutes. The cells were washed and 100 μL of 4% paraformaldehyde was added to the cells. The cells were stored in paraformaldehyde until flow cytometry was performed.

### Immunofluorescence Staining

A glass bottom black sided 96 well plate (Greiner Bio-one) was coated with RTC1 and allowed to incubate for at least 4 hours. The wells were washed with PBS twice. Cells were seeded onto the plate at a concentration of 2,000 cells/well. The cells were incubated at 37°C with 5% CO2 overnight. Then the cells were fixed with 3.7% Formaldehyde for 15-20 minutes, the membranes of the cells were lysed with 0.1% Triton-X for 10 minutes, and finally the cells were blocked with 1% BSA for 1 hour. The cells were incubated in a 1:250 dilution of Von Willebrand Factor Antibody (vWF; Abcam) for 1 hour. Then a AlexaFlour 488 Goat anti-rabbit secondary antibody (Abcam) at a 1:200 dilution and a Rhodamine conjugated *Ulex Europaeus Agglutinin 1* (UEA-1) at a concentration of 10 μg/mL were added and allowed to incubate for 30 minutes. Finally, the cells were stained with 300 nM DAPI (Thermo Fisher) for 3 minutes. The cells were washed 3 times with PBS between steps and after the DAPI staining. The cells were then imaged on the Nikon AX-R confocal microscope.

### Tube Formation Assays

15 well µ-angiogenesis plates (ibidi) were placed into a -80°C freezer and allowed to cool for at least 2 hours. On ice, 10 μL Matrigel was added to each well. The plates were added to a petri dish with 1-2 mL of PBS and placed into an incubator at 37°C with 5% CO2. After 1-3 hours of incubation, the 50 μL of cell suspension at 80,000 cells/mL were added to each well. The cells were imaged using the Lionheart FX Automated Microscope (BioTek Instruments) every hour for 12 hours. The angiogenic potential of each well was determined by using a free ImageJ plugin in called Kinetic Analysis Vasculogenesis (KAV)^36^.

### Proliferation Assays

All the plates mentioned in this section were coated with RTC1 the day before and allowed to incubate overnight at 37°C with 5% CO2. The plates were washed twice before the cells were added.

#### Label-Free

Cells were seeded in a 96 well flat-bottom tissue culture plate (VWR, Radnor, PA) at 6,000 cells per well in 250 μL of EGM2 and were incubated at 37°C with 5% CO2. After 2 hours, cells underwent a high contrast cell counting protocol developed by BioTek Instruments on a Lionheart FX Automated Microscope (BioTek Instruments). Cells were counted based on images taken every 2 hours for 48 hours.

#### WST-1

Cells were seeded in a 96 well flat-bottom tissue culture plate (VWR) at 6,000 cells per well in 250 μL of EGM2 and were incubated at 37°C with 5% CO2 overnight. 10 µL of WST-1 (Sigma Aldrich) was incubated for 2 hours and put on a shaker plate for 1 minute. Then readings were taken on a microplate reader (Spark).

#### Cell Cycle Analysis

Cells were grown to 70%-80% confluency in T75 flasks (VWR). They were then treated with serum free media to try to have the cells at a similar point proliferatively. Cells were stained with Click-iT EdU and PI/RNase using established protocols from Thermo Fisher. The fluorescence of both stains was quantified using Flow Cytometry and analyzed using FlowJo (FlowJo, LLC.).

### Wound Healing Assay

A 24 well flat-bottom tissue culture plate (VWR) was coated with RTC1 and allowed to incubate for at least 4 hours. The wells were washed with PBS twice and then were allowed to dry until the 2 well inserts (ibidi) can adhere. 70 μL of cell suspension at 80,000 cells/mL were added to each side of the insert and 300 μL of EGM2 was added to the surround well. The plate was incubated overnight. The inserts were removed, the media was aspirated, and 500 μL of EGM2 was added to the well. The cells were imaged using the Lionheart FX Automated Microscope (BioTek Instruments) every hour for 24 hours. The area calculations were based on a protocol developed by BioTek Instruments.

### Senescence

Cells were grown up to passage 11. They were then seeded into a 96 well flat-bottom tissue culture plate (VWR) that had been RTC1 coated at 1000 cells/well. The cells were incubated at 37°C with 5% CO2 overnight. The cells were then fixed and stained with a Senescence Detection Kit (Abcam) as per manufacturer’s protocol. The cells remained in the solution for 20 hours and then the number of blue cells were counted.

### Cytokine Secretion

Supernatant was collected from cells at approximately 70-90% confluency. The supernatant was filtered through a 0.2 μm filter to remove dead cells. Proteomics were done using an established protocol from R&D Systems until the final steps. The cards were imaged using a ChemiDoc system. The angiogenesis associated proteins were found using a Proteome Profiler Human Angiogenesis Array Kit (R&D Systems) and the inflammation associated proteins were found using Proteome Profiler Human Cytokine Array Kit (R&D Systems).

### Real-time qPCR

ECFCs and PRECs were grown on RTC1 coated tissue culture 6 well plate (VWR). There were four biological replicates selected for preeclamptic and healthy ECFCs (n=3). 2 wells were pooled for each biological replicate. Real-time qPCR was performed as previously described^37^. Gene expression assays for *LYVE-1, Prox-1, VEGFR3,* and *GAPDH* were used in addition to the TaqMan™ Array Human Angiogenesis Panel (Thermo Fisher). Three biological replicates selected for preeclamptic and healthy ECFCs (n=3).

### Cell Treatment

ECFCs and PRECs were treated with 5 nM Fc Chimera (R&D Systems) or 100 ug/mL NRP-1 Antibody (R&D Systems) every other day for 5 days.

### Statistical Analysis

Analysis was done with GraphPad Prism 10 (GraphPad Software Inc.). Student’s t-test or ANOVA was used to compare ECFCs and PRECs. Statistical significance was set at ****p<0.0001, ***p<0.001, **p<0.01, and *p<0.05.

## Results

### Preeclampsia Does Not Alter Endothelial Marker Expression

To establish that PRECs maintain endothelial marker expression, comparative flow cytometry was performed for common endothelial markers CD31 (PECAM-1), CD34, and CD144 (VE-Cad) **(Figure 1A,C)**. There was no significant expression found between the cell lines **(Figure 1B,D).** While there was some slight variability in expression of CD31 between biological replicates, it was not statistically significant (P = 0.4035). ECFCs and PRECs were also stained for vWF and UEA-1 and expressed similar patterns **(Figure 1E)**. Collectively, these results indicate that preeclampsia is not affecting endothelial marker expression in ECFCs.

**Figure 1.**
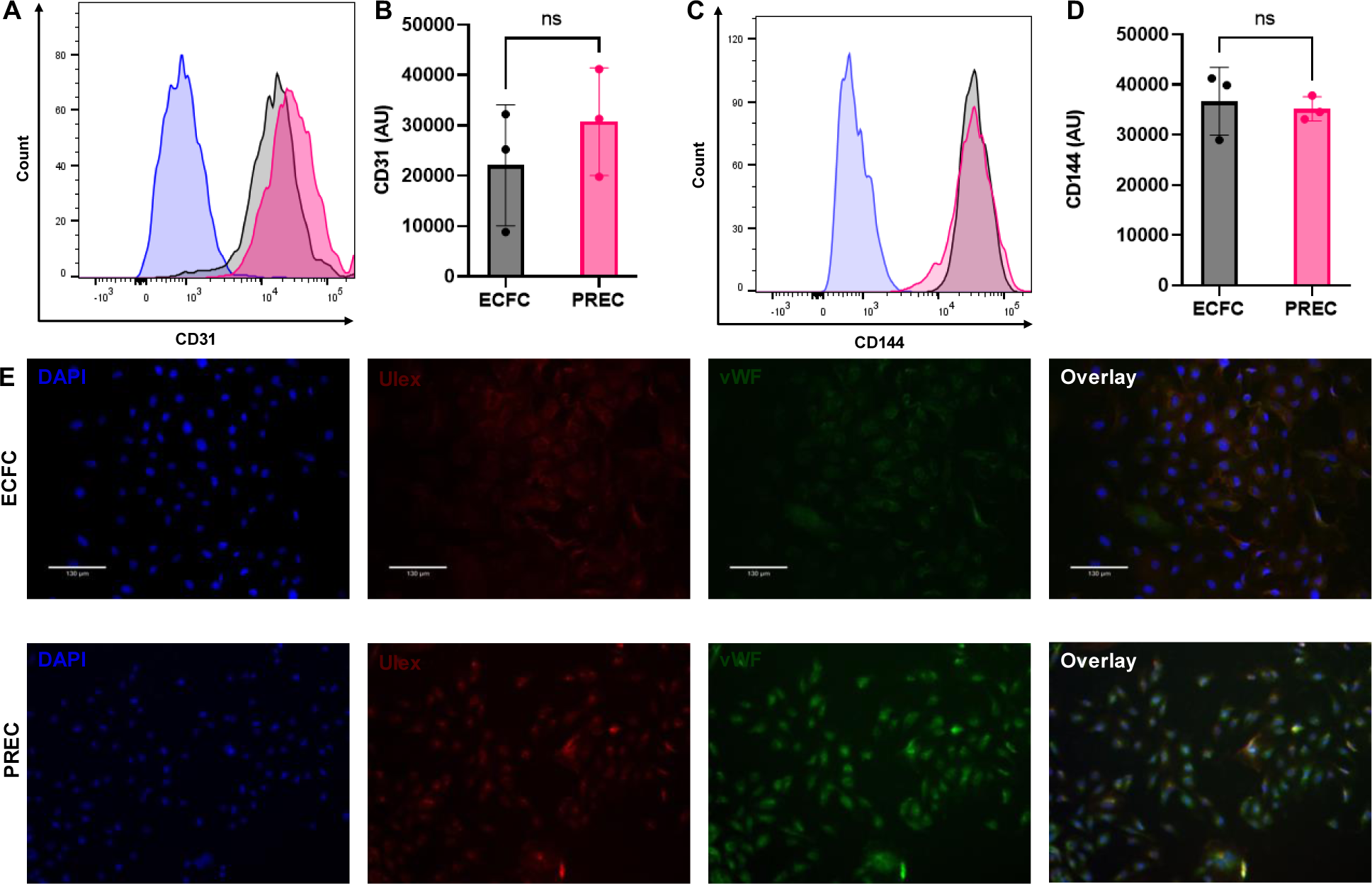
Expression of endothelial markers (A) CD31 and (C) CD144 were measured in both ECFCs and PRECs. Control is shown in blue, the ECFCs are in black, and ECFCs are in pink. Their mean values were quantified (B&D). (E) Ulex (Red) and van Willebrand’s Factor (vWF; green) were stained alongside DAPI (blue). Scale bar = 130 μm.

### PRECs have Decreased Tube Forming Ability

Since preeclampsia is known to cause endothelial dysfunction, the first characteristic looked at was PRECs’ ability to form tubes. ECFCs and PRECs were allowed to form tubes on Matrigel for 9 hours. Then the networks were imaged and quantified using a free ImageJ plugin called kinetic analysis vasculogenesis (KAV)^34^. The ECFCs formed a more organized network with multiple complete loops visible **(Figure 2A)**, while the PRECs were more dysregulated **(Figure 2B)**. Although KAV produces 10 parameters to quantify the tube network, network size and the average tube length were looked at to confirm changes in angiogenic potential. Network size shows the area within the loops and shifts in this parameter indicate morphological changes. PRECs saw a decrease in network size as the loops tended to be smaller than their healthy counterparts **(Figure 2C)**. When the networks branch and split, the tube length tends to be shorter. The PRECs tend to have less completed loops and less branching causing them to have longer tubes **(Figure 2D)**.

**Figure 2.**
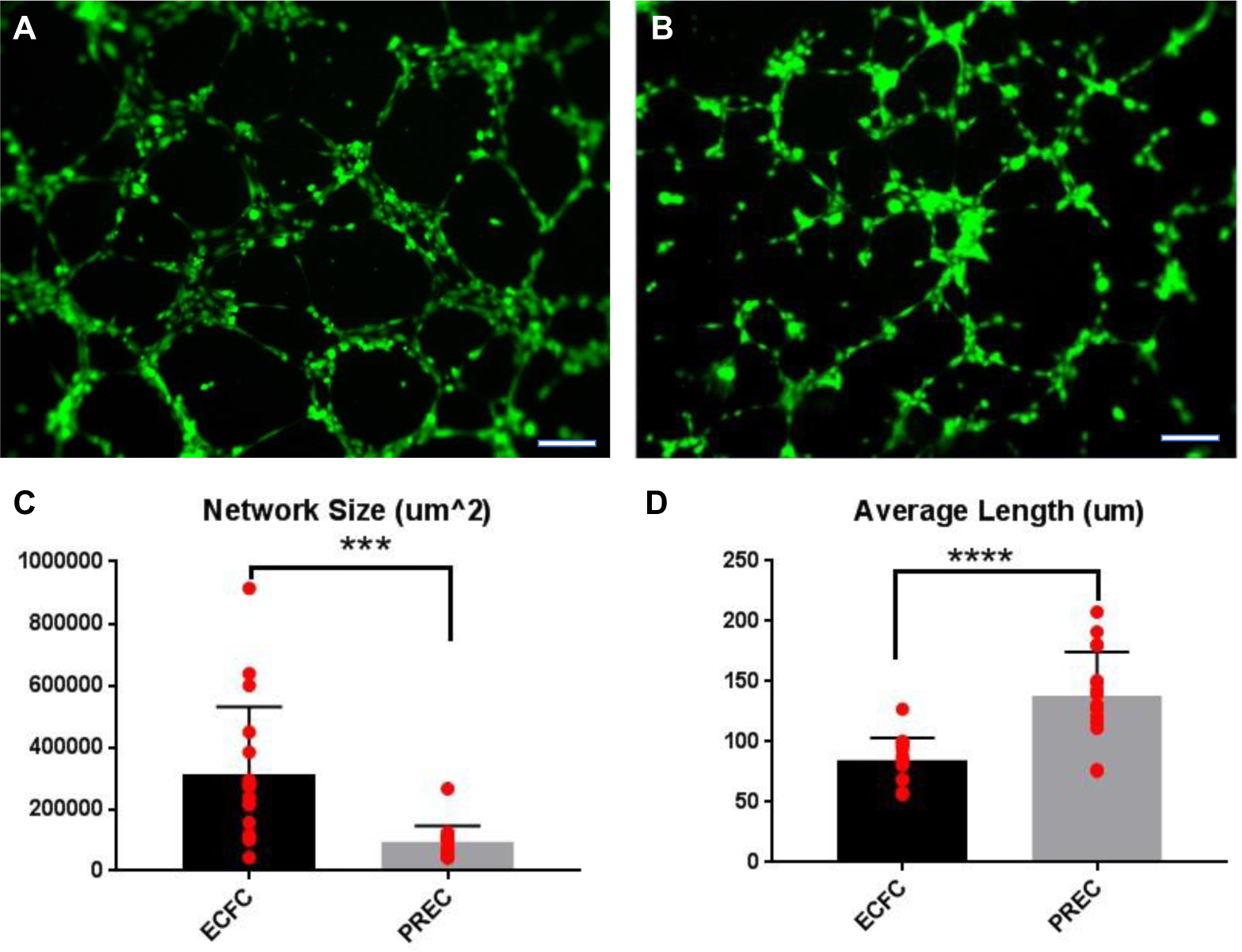
Angiogenic potential of ECFCs was measured using a tube formation assay. The assay was performed with (a) ECFCs and (B) PRECS. Kinetic analysis Vasculogenesis (KAV) was used to quantify the (C) network area and (D) average tube length. ***p < 0.001, **** p < 0.0001. Scale bar = 250 μm.

### Preeclampsia Causes Functional Differences in ECFCs

To determine the cause of this decreased tube forming ability, other functional characteristics were assessed. The first being proliferation. Using a protocol developed by Biotek Instruments, high contrast light was used to count cells without the addition of a label **(Figure 3A)**. By 40 hours, there was a statistically significant difference between the number of PRECs compared to the number of ECFCs per well. Since this method is less established, two other proliferation assays were performed. WST-1, which indicates proliferation based on mitochondrial activity, saw a slight increase in PREC proliferation **(Figure 3B)**. However, it was not statistically significant as there is high variability between biological replicates since the different samples came from different donors. Cell cycle analysis was performed using PI/RNase for DNA content and Click-iT EdU to indicate the S phase **(Figure 3C)**. PRECs had a lower percentage of cells in the G0 and G1 phases and an increased proportion of cells in the S phase **(Figure 3D)**. The percentage of G2 and M phase cells was similar between ECFCs and PRECs. Collectively, these results indicate that the PRECs proliferate more rapidly than the ECFCs, an already highly proliferative cell type^27^.

**Figure 3.**
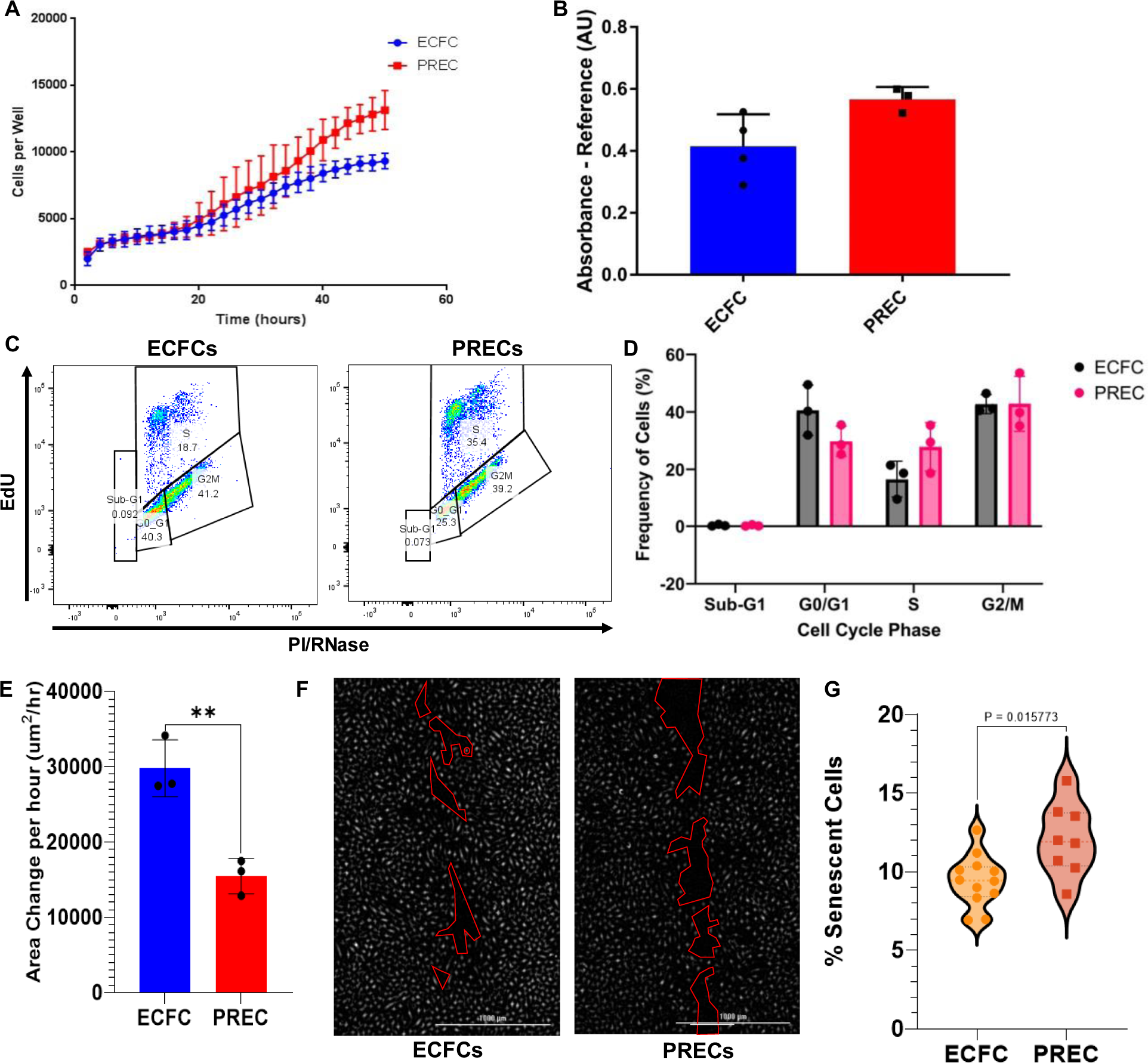
Functionality was measured in ECFCs and PRECs using various assays. Proliferation was examined using (A) a label-free proliferation assay, (B) WST-1, and (C-D) cell cycle analysis. (E) A wound healing assay was used to quantify migration. **p<0.01. After 12 hours, (F) the wound for the ECFCs and PRECs were imaged. (G) Senescence was investigated using a β-gal assay. Scale bar = 1000 μm

*In utero* conditions such as gestational diabetes mellitus (GDM) have been shown to affect the ability of cells to migrate^31^. Therefore, it was predicted that preeclampsia may also have an effect. Using a wound healing assay with ibidi 2 well inserts both ECFCs and PRECs were tested. It was seen that the ECFCs closed the gap quicker than the PRECs **(Figure 3E)**. By 12 hours, there was a noticeably larger gap in the PRECs compared to the ECFCs **(Figure 3F)**.

Finally, a β-gal senescence assay was performed. The cells were incubated overnight in the staining solution (20 hours) and then imaged. The percentage of senescent cells seen in PRECs was increased compared to ECFCs **(Figure 3G)**. Collectively, these findings indicate that preeclampsia affects the functionality of the ECFCs making the cells more proliferative and senescent, while decreasing migration.

### Preeclampsia Dysregulates Angiogenic and Inflammatory Cytokine Production

Based on these findings, possible causes of these functional changes were further investigated. The first aspect looked at was cytokine production. Since preeclampsia is a hypoxic disease, the PRECs may see increased inflammatory cytokine production. Proteomics analysis was performed on the supernatant from ECFCs and PRECs. We found that there was an increase in CXCL1/GROa, IL-8, and MIF in PRECs though it was not significant. However, ECFCs saw a higher amount of Serpin E1 expression than their preeclamptic counterparts **(Figure 4A)**. When proteomics analysis was done for angiogenesis associated proteins, a major dysregulation was seen in PRECs. However, there was not a clear trend **(Figure 4B)**. Serpin E1 and IL-8 were found to follow a similar pattern to what was seen in the inflammatory panel. In addition, urokinase plasminogen activator (uPA), which is inhibited by Serpin E1, also follows the expected trend, and decreases in PRECs.

**Figure 4.**
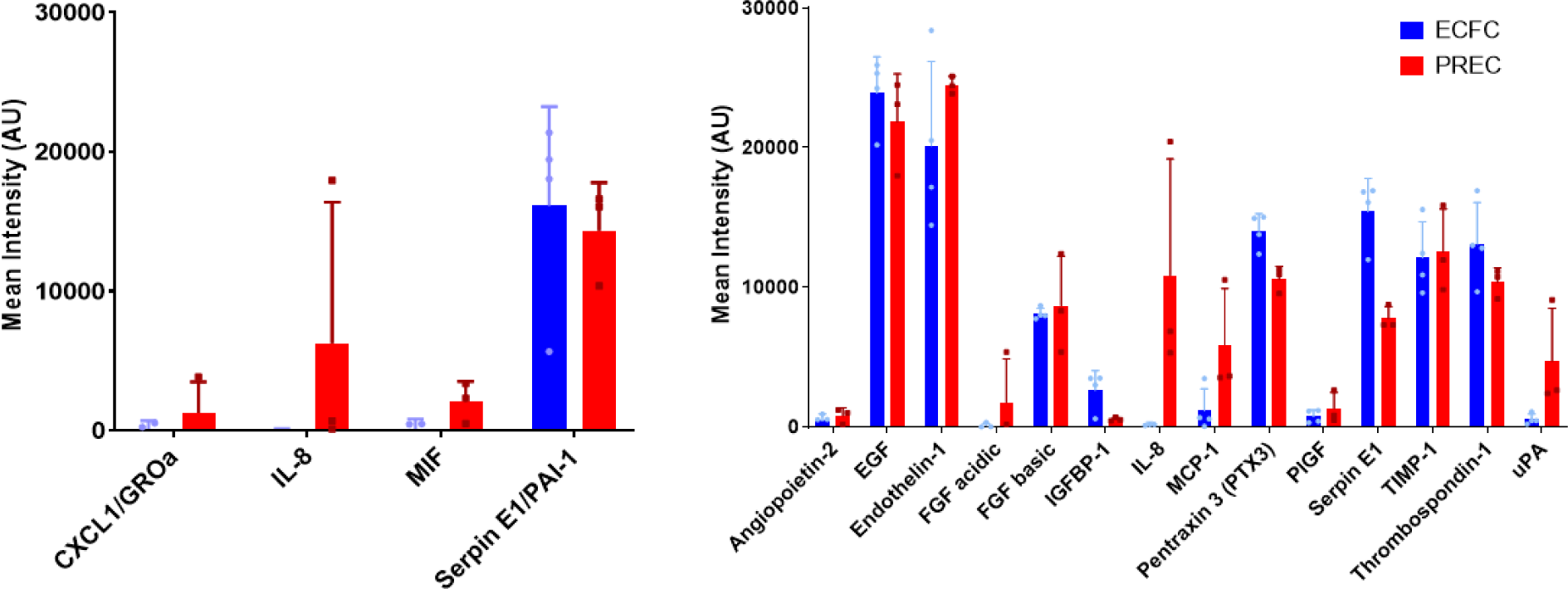
Proteomics profiles for **(A)** Inflammatory and **(B)** Angiogenesis associated proteins.

### PRECs Express Dysregulated Angiogenesis Associated Genes

After observing the functional changes in PRECs, the genetics were looked at. A 94 gene angiogenesis PCR panel (Thermo Fisher) was run on 4 ECFC and 4 PREC cell lines **(Figure 5A)**. It was found that Fibulin 5 (*FBLN5*) and proto-oncogene, receptor tyrosine kinase (*KIT*) were significantly dysregulated and had a fold change greater than 2 (**Figure 5B,C)**. Fibulin 5 has been shown to decrease endothelial sprouting, proliferation, and invasion in other types of endothelial cells^38,39^. KIT can influence many cellular functions including migration and tube formation^40^. Hypoxia, which is commonly associated with preeclampsia, has been shown to upregulate both Fibulin 5 and KIT^41,42^.

**Figure 5.**
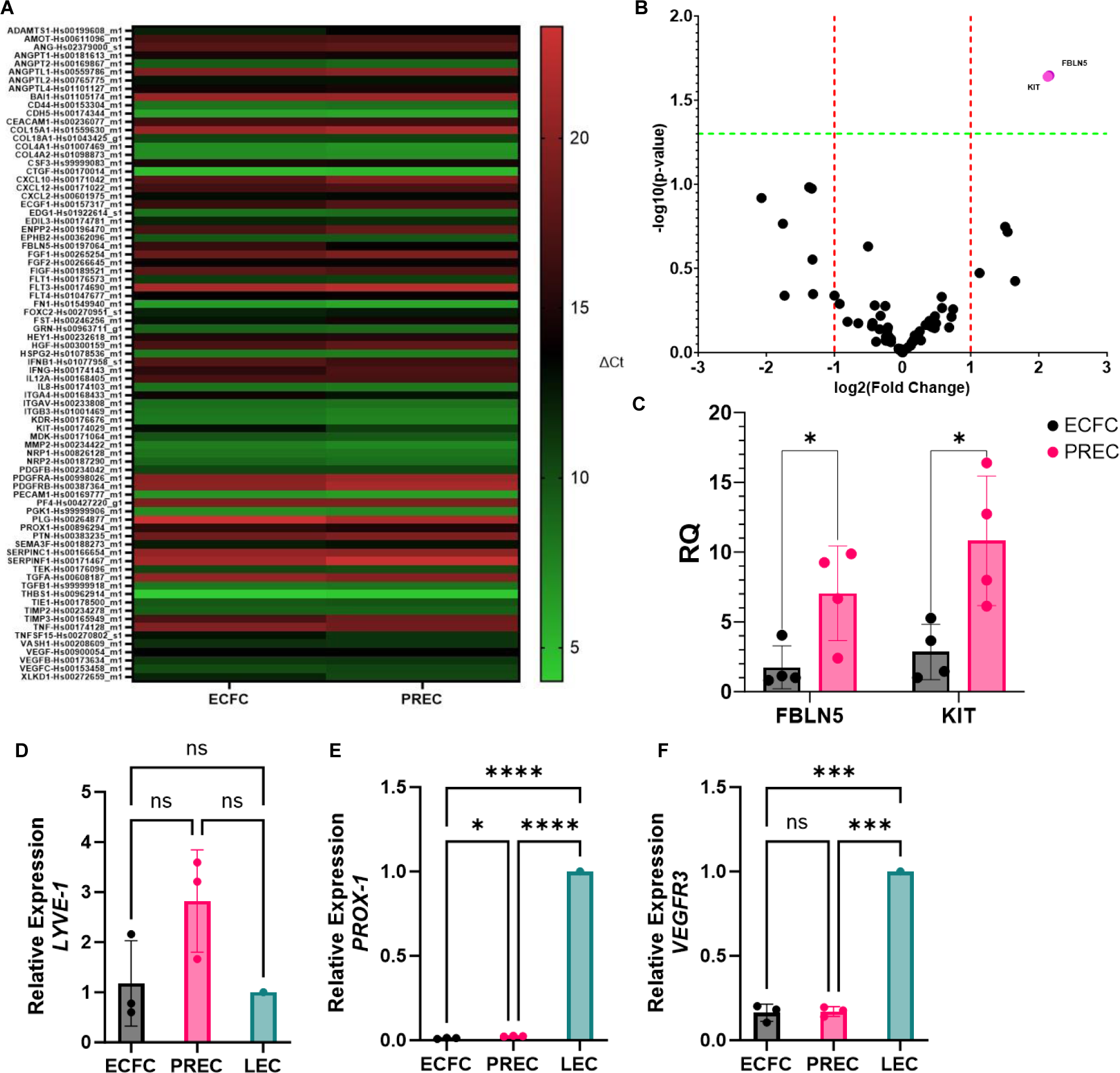
Real-time qPCR. **(A)** The Taqman Angiogenesis Panel was run for 94 genes and 2 endogenous controls (GAPDH and 18S). Three biological replicates were used for PRECs and ECFCs (n=3). **(B)** A volcano plot from the data from A. **(C)** Fold-change for Fibulin 5 and KIT. PCR data for lymphatic genes **(D)** *LYVE-1,* **(E)** *Prox-1,* and **(F)** *VEGFR3*. GAPDH is used as endogenous control. Three biological replicates were used for PRECs, ECFCs, and LECs (n=3). ****p<0.0001, ***p<0.001, and *p<0.05

Studies have shown an increase in lymphatic mimicry in endothelial cells during pregnancy^43^. To determine if lymphatic genes are dysregulated in preeclampsia, PCR was run for lymphatic vessel endothelial hyaluronan receptor 1 *(LYVE-1*), Prospero homeobox protein 1 *(Prox-1),* and Vascular Endothelial Growth Factor Receptor 3 (*VEGFR3*) compared to lymphatic endothelial cells (LECs), where they are natively expressed. *VEGFR3* was found to not be significantly different between ECFCs and PRECs. However, both groups had lower expressions than LECs **(Figure 5F)**. PRECs appear to have increased *LYVE-1* expression compared to both LECs and ECFCs. Though due to high variability between biological samples, it is not statistically significant **(Figure 5D)**. *Prox-1* is significantly upregulated in PRECs compared to ECFCs but is lower than what is seen in LECs **(Figure 5E)**. Overall, there are slight shifts in the lymphatic marker expression in PRECs, however, it needs to be investigated further.

### Neuropilin-1 Improves the Angiogenic Potential of PRECs

Neuropilin-1 (NRP-1), a co-receptor of Vascular Endothelial Growth Factor Receptor 2 (VEGFR2), has been shown to effect both proliferation and senescence in ECFCs^44^. To determine if this co-receptor was affected by preeclampsia, plays a role in the functional dysregulation found in ECFCs, comparative flow cytometry was performed **(Figure 6A)**. It was observed that there was no significant shift in NRP-1 expression between ECFCs and PRECs. However, the PRECs had higher variability between biological replicates, indicating an inconsistent dysregulation. The cells were then treated 5 nM Fc Chimera (R&D Systems) or 100 ug/mL NRP-1 Antibody (R&D Systems) every other day for 5 days. The cells were then seeded onto Matrigel and allowed to form tubes **(Figure 6B)**. Using KAV for quantification, it was found that the Fc treatment improved angiogenic potential. The Fc was found to increase the number of closed networks **(Figure 6C)** and increase the average tube length **(Figure 6D)** to be similar to the untreated ECFCs. The antibody appeared to decrease both of these parameters **(Figures 6C,D)**. Due to the highly variable expression of NRP-1 in PRECs, NRP-1 treatment needs to be investigated further to determine if the Fc can improve functionality consistently.

**Figure 6.**
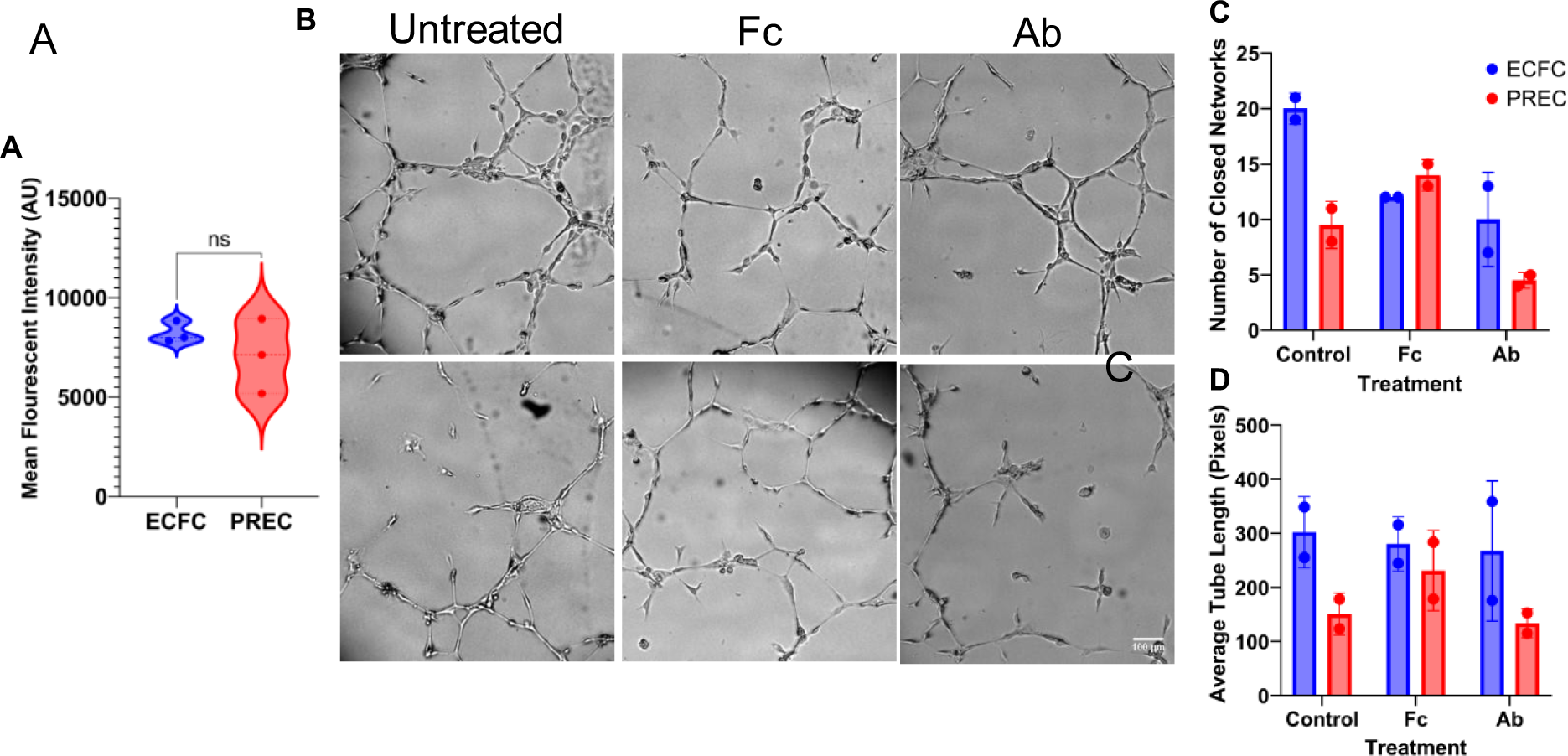
(A) NRP-1 expression was quantified based on mean fluorescent intensity. (B) Tube formation done with an activator (Fc) and inhibitor (Ab) of NRP-1. (C) The number of closed networks and (D) average tube length were quantified using KAV. Scale bar = 100 μm

## Discussion

As a result of preeclampsia, PRECs exhibit decreased migratory and tube forming abilities as well as increased proliferation. There is a balance needed between proliferation and migration to have successful vessel development. Computational models have shown that high proliferation and low migration cause less sprouting and more twisted vasculature^45^. In addition, the amount of senescence increased in PRECs. This increase in proliferation may be an overcompensation for premature senescence. A better understanding of these functions in relation to each other is important for understanding how preeclampsia is affecting the ECFCs.

In addition to functional dysregulation, protein production is also disrupted. The inflammatory cytokines CXCL1/GROa, IL-8, and MIF were all increased while Serpin E1 was decreased in PRECs. As preeclampsia associated with hypoxia, the increase in these cytokines follows expected trends. In addition, many angiogenesis associated factors decreased in PRECs, including Angiopoietin-2, TIMP-1, and Pentraxin 3, while only a couple of factors increased (e.g., MCP-1, PlGF, and Prolactin) compared to ECFCs. Additionally, Serpin E1 follows the same pattern as seen in the PRECs’ inflammatory cytokine profile. The protein inhibited by Serpin E1, uPA, increased as expected. As angiogenic dysregulation is a known characteristic of preeclampsia, this dysregulation is expected. However, highlighting proteins like Serpin E1, IL-8, and uPA, which are noticeably affected by this disorder, can give insight into possible targets for treatment or potential biomarkers to better understand preeclampsia.

On the genetics side, the two genes that were found to be significantly upregulated in PRECs were *KIT* and *FBLN5*. *KIT* is important for the identification of progenitor cells as well as migration, differentiation, and capillary tube formation^40^. Generally, overexpression leads to increased angiogenesis and proliferation^42^. *FBLN5* is an extracellular matrix glycoprotein that plays a role in endothelial cell attachment and proliferation. Overexpression of *FBLN5* has been found to increase proliferation, increase apoptotic markers in matured endothelial cells, and decrease angiogenic potential as it attenuates Angiopoietin-1 and VEGF signaling pathways^38,39,46^. Together, these genes appear to play a role in the change in angiogenic potential seen in previous sections. However, further research is needed to determine the extent of their effect.

Lymphatic mimicry has been shown to occur during pregnancy. LYVE-1, Prox-1, and VEGFR3 were all expressed in spiral arteries during remodeling^43^. These markers are also expressed in other parts of the placenta. LYVE-1 has been shown to be expressed over CD44, a hyaluronic acid receptor that is a homolog of LYVE-1, in the trophoblasts and villous endothelial cells^47^. *Prox-1* was also seen to be expressed in these cells at the mRNA level, but not the protein level. In addition, abnormal lymphatic development in the decidua has been observed^48^. Despite these markers being observed during pregnancy, lymphatic mimicry has not been looked at in ECFCs during preeclampsia. This study looked at *LYVE-1*, *Prox-1*, and *VEGFR3* expression in ECFCs and PRECs. *LYVE-1*, despite not being significant due to variability between clinical samples, had higher expression (average 2.8 fold change) in PRECs compared to their healthy counterparts and LECs. *LYVE-1* expression in ECFCs is increased compared to mature endothelial cells like HUVECs^49^. Therefore, the lack of significance between the ECFCs and LECs is not unexpected. Prox-1 as the master regulator of lymphatic markers was expected to be upregulated in PRECs as we see an increase in other lymphatic markers. However, this upregulation is still significantly lower than what was observed in native LECs. VEGFR3 was significantly lower in ECFCs and PRECs than LECs. Together, these findings indicate that ECFCs are experiencing a minor upregulation of lymphatic markers, but the expression is still lower than lymphatic tissues. A comparison with other cells that express lymphatic markers during pregnancy like trophoblasts or spiral artery cells may provide insight into the extent of lymphatic mimicry seen in ECFCs and PRECs.

NRP-1 treatment has been shown to affect proliferation and senescence in ECFCs^44^. NRP-1 expression is highly variable in PRECs, which makes it difficult to determine the effectiveness of the treatment. Agonists (Fc) and antagonists (Ab) were used to treat ECFCs and PRECs. Since the majority of PREC cell lines in this study expressed lower amounts of NRP-1, the addition of the agonist appears to improve the tube forming ability. However, testing with more biological replicates is needed to verify this result and determine if a decrease in NRP-1 expression is the expected trend.

Collectively, these results show a shift in the genetic and functional profiles of ECFCs in preeclampsia. Some of these abnormally expressed proteins could be investigated further as potential biomarkers. In addition, treatment of NRP-1 can improve tube forming ability. Findings from this study highlight proteins and genes in ECFCs affected by preeclampsia that could be studied and treated in future experiments.

## Abbreviations

ECFCs: Endothelial Colony-Forming Cells
EPCs: Endothelial Progenitor Cells
GDM: Gestational Diabetes Mellitus
HUVECs: Human umbilical vein endothelial cells
KAV: Kinetic Analysis Vasculogenesis
LECs: Lymphatic endothelial cells
LYVE-1: Lymphatic vessel endothelial hyaluronan receptor 1
PlGF: Placental Growth Factor
PRECs: Endothelial Colony-Forming Cells isolated from a preeclamptic pregnancy
Prox-1: Prospero homeobox protein 1
sEng: Soluble endoglin
sFlt-1: Soluble fms-like tyrosine kinase
VEGF: Vascular endothelial growth factor
VEGFR3: Vascular Endothelial Growth Factor Receptor 3

## Acknowledgements

We acknowledge support from the University of Notre Dame through “Advancing Our Vision” Initiative in Stem Cell Research and Scientific Wellness Initiative, Harper Cancer Research Institute – American Cancer Society Institutional Research Grant (IRG-17-182-04), American Heart Association through Career Development Award (19-CDA-34630012 to D.H.-P.), and from National Institutes of Health (R01-HL-094725 to L.S.H. and 1R35-GM-143055-01 to D.H.-P.). We would like to thank the Notre Dame Bioinformatic Core and AngioBioCore at Indiana University Simon Comprehensive Cancer Center (NIDDK/NIH U54-DK-106846 and P30 CA-082709) for isolating and characterizing cord blood samples. This publication was made possible, with support from the Indiana Clinical and Translational Science Institute (I-CTSI) funded, in part by Grant Number ULITR001108 from the NIH for Advancing Translational Sciences, Clinical and Translational Science Awards.

## Author contributions

E.H., L.A., D.H., L.S.H., and D.H.-P. conceived the ideas, designed the experiments, interpreted the data, and wrote the manuscript. E.H., L.A., S.S., E.J., E.N. conducted the experiments and analyzed the data. L.S.H and D.H.-P. supervised the study. All authors have approved of the manuscript.

## Data and materials availability

Additional material and data which contributed to this study are present in the **Supplementary Information**.

