## Supplemental Table 1 for "The Effects of Preeclamptic Milieu on Cord Blood Derived Endothelial Colony-Forming Cells"

**Supplementary Information**

Table S1. Antibody and Treatment Reagents

| Reagent | Manufacturer | Part Number | Dilution/Final Concentration |
|---|---|---|---|
| CD31 Antibody (Flow) | R&D Systems | FAB3567A | 10 μL/$10^6$ cells |
| CD34 Antibody (Flow) | Abcam | ab187284 | 1 μg/mL |
| CD144 Antibody (Flow) | BD Pharmingen | 560410 | 20 μL/test |
| vWF Antibody | Abcam | ab154193 | 1:250 |
| Alexa Flour 488 Secondary Antibody | Abcam | ab150077 | 1:200 |
| UEA-1 Antibody | Vector Labs | RL-1062 | 10 μg/mL |
| DAPI | Thermo Fisher | D1306 | 300 nM |
| NRP-1 Antibody (Flow) | R&D Systems | FAB3870A | 10 μL/$10^6$ cells |
| NRP-1 Fc Chimera | R&D Systems | 566-NNS-025 | 5 nM |
| NRP-1 Antibody | R&D Systems | AF3870-SP | 100 ug/mL |
